## Supplementary Figures & Note for "Symmetry-adapted Markov state models of closing, opening, and desensitizing in α7 nicotinic acetylcholine receptors"

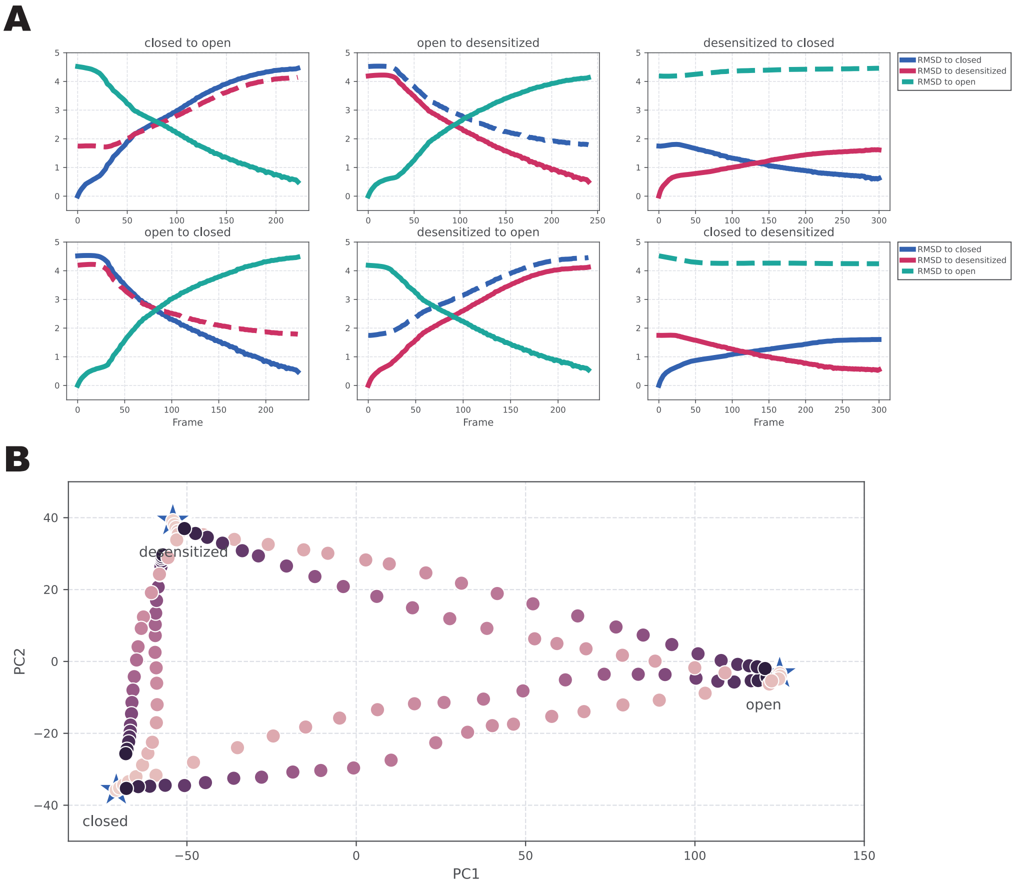


**Figure S1. Interpolation analysis of Climber trajectories.** (**A**) Root mean square deviation (RMSD) of each interpolation from the three structural models. (**B**) Projection of selected seeds along each interpolation onto the principal components generated from the structural models.

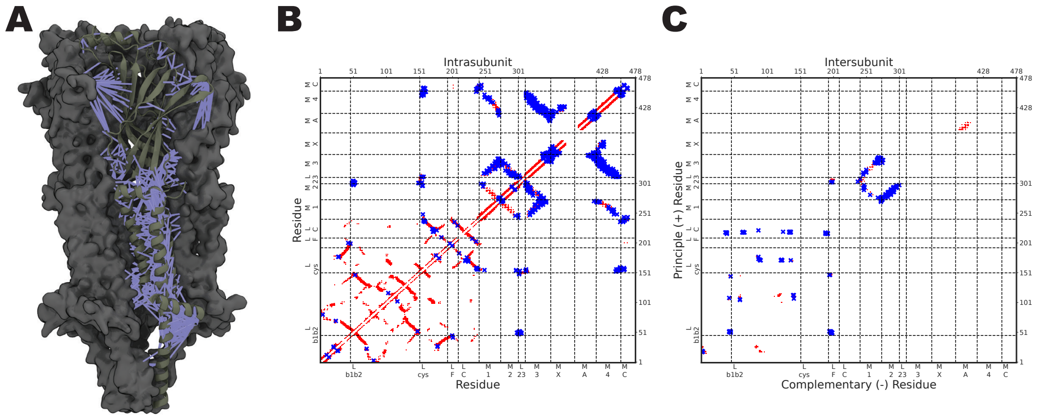


**Figure S2. Feature selection and contact analysis.** (**A**) The selected features for the analysis are highlighted in blue, representing interatomic distances. (**B**) Intrasubunit $C_\alpha$ contacts are shown in red, while the selected contacts are highlighted in blue. (**C**) Intersubunit $C_\alpha$ contacts are shown in red, and selected contacts highlighted in blue.

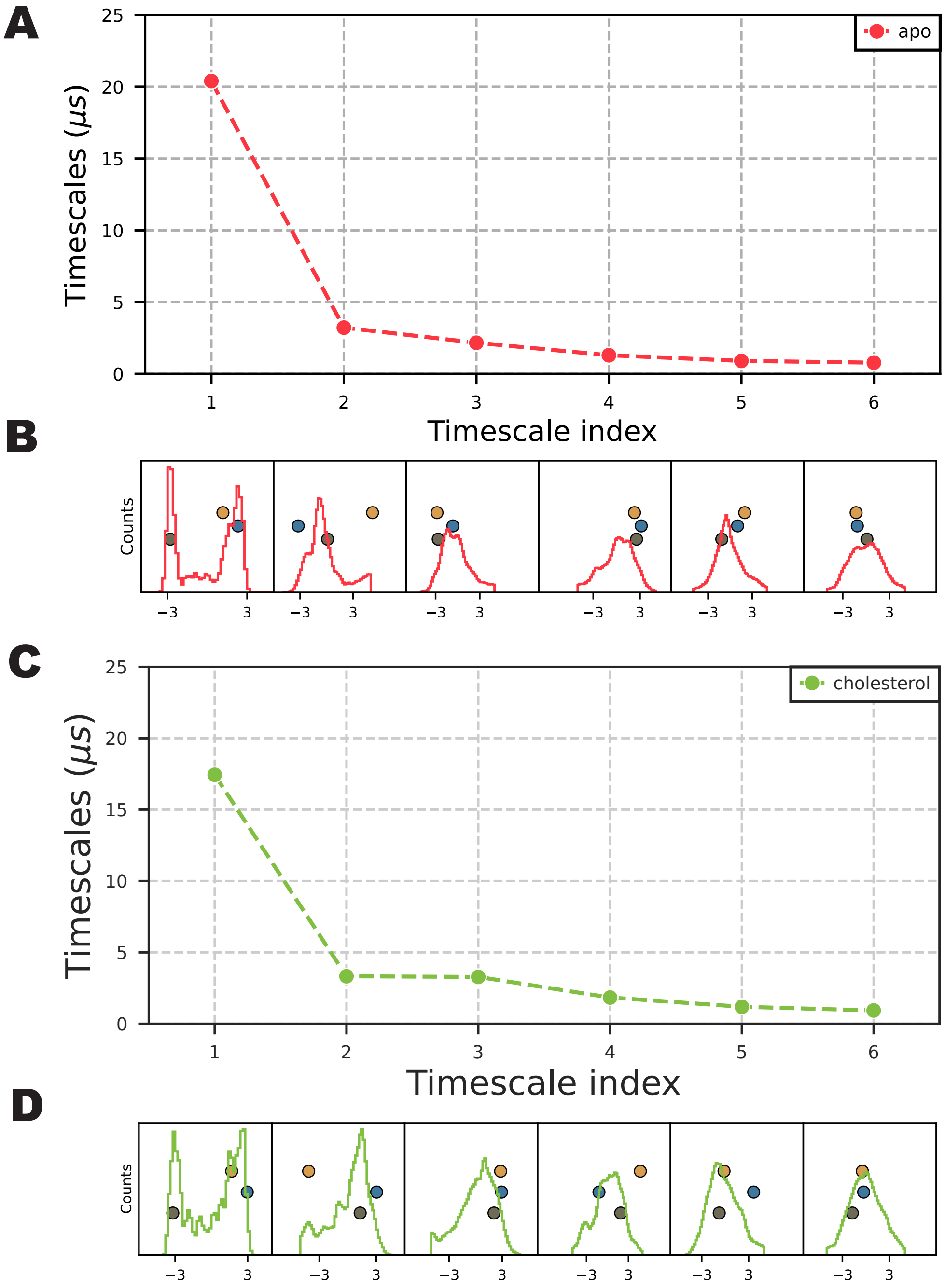


**Figure S3. Implied timescale of each independent components.** (**A, C**) Implied timescale of each independent component for the apo and CHOL systems. (**B, D**) Histogram of each independent component for the apo and CHOL system. Structural models are projected onto the plot (grey: closed; blue: open; yellow: desensitized).

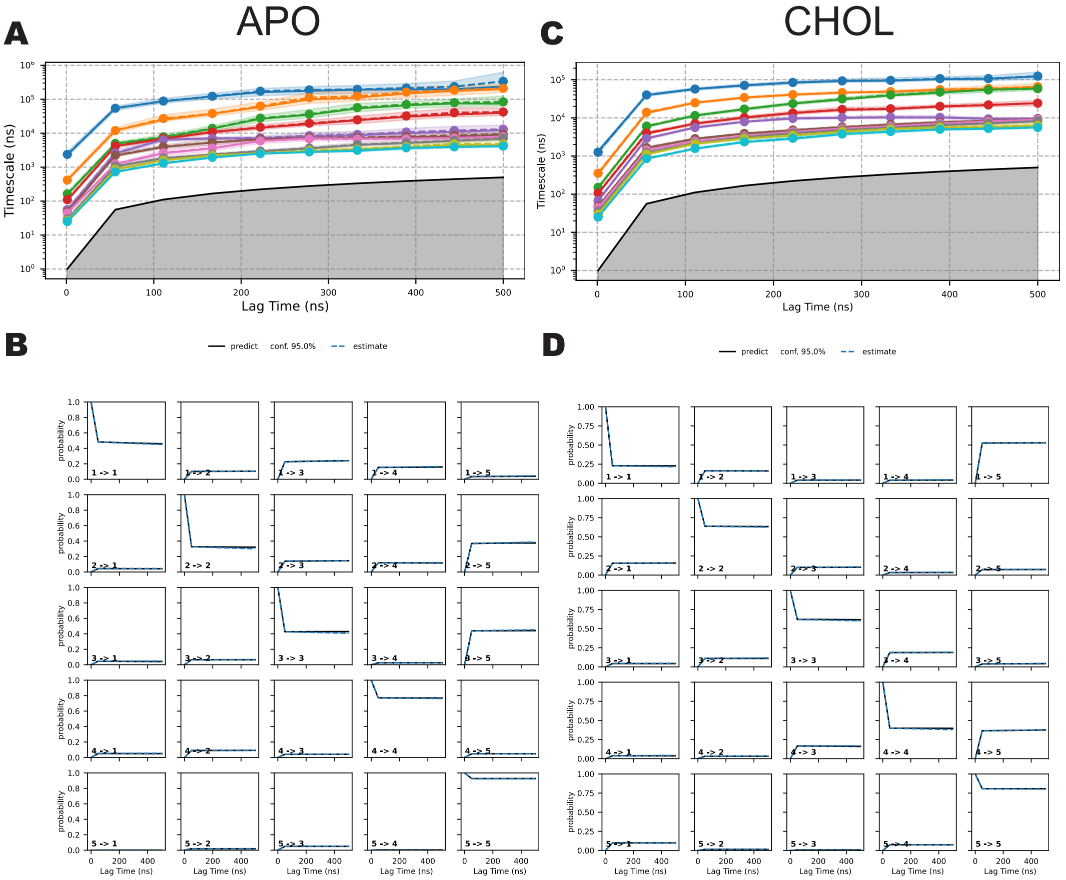


**Figure S4. Assessment of convergence of the MSMs.** (**A, C**) Implied timescale with MSMs estimated with different lag times for apo and CHOL systems. (**B, D**) Chapman-Kolmogorov tests with five states for apo and CHOL systems.

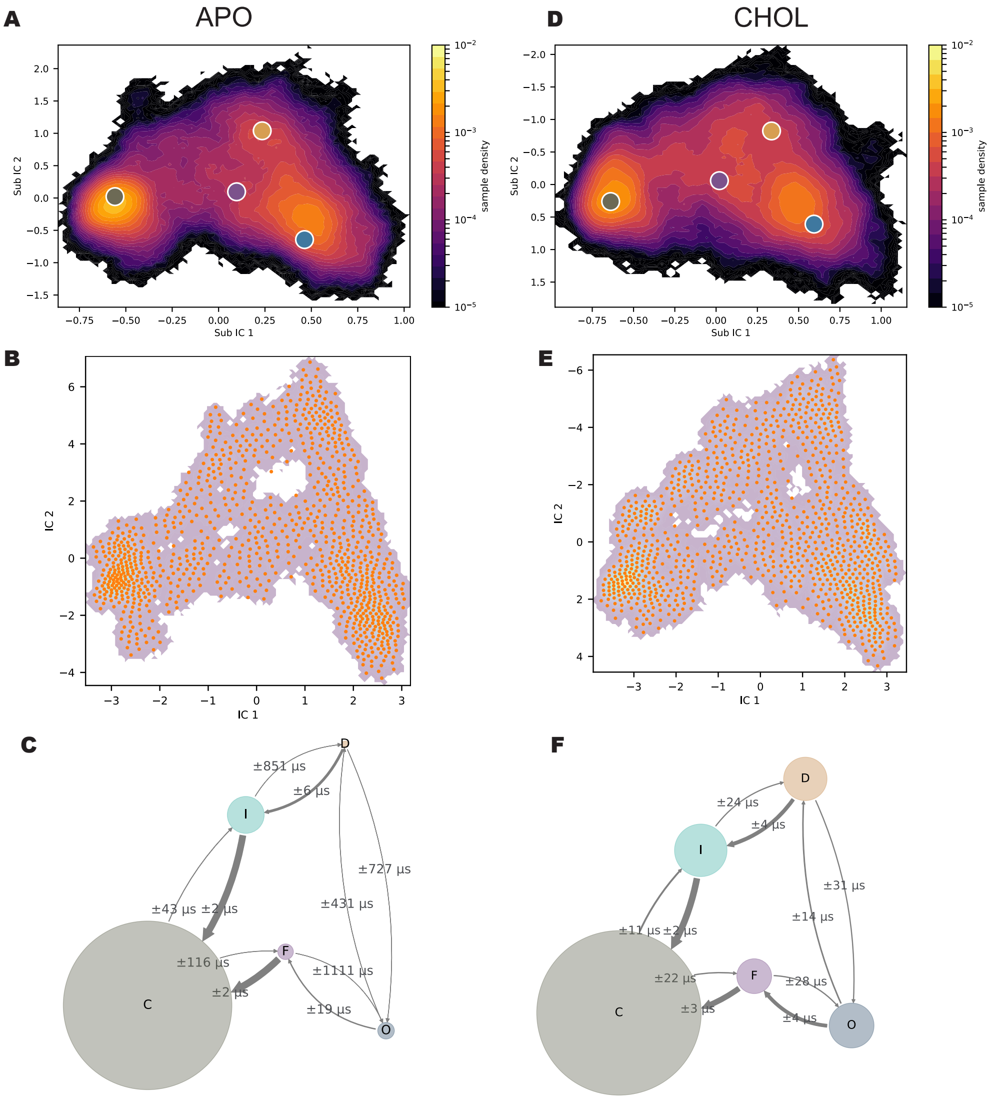


**Figure S5. Microstates in MSMs and kinetic error estimation.** (**A D**) Projection of the four structural models onto sub IC1-IC2 coordinate density map for apo and CHOL systems. (**B E**) 1000 microstates assigned with k-mean clustering for apo and CHOL systems. (**C F**) The error estimation for each mean first passage time for apo and CHOL systems.

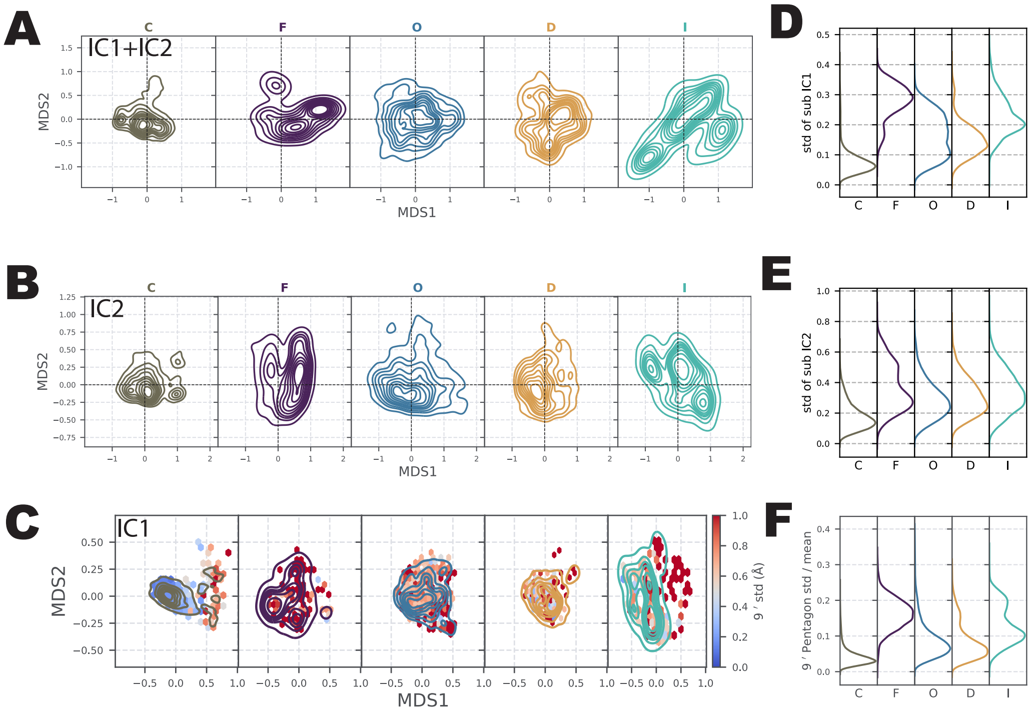


**Figure S6. Standard deviation and MDS of other ICs.**(**A**) Projection of sub independent components (IC1+IC2) onto MDS space. (**B**) Projection of sub IC2 onto MDS space. (**C**) Projection of sub IC1 onto MDS space. The histogram of the standard deviation of the five edges on the 9′ $C_\alpha$ pentagon is plotted as a heatmap. (**D,E,F**) Distribution of the standard deviation of sub IC1, IC2, and normalized 9′ $C_\alpha$ pentagon edge lengths in each macrostate. All distributions are weighted by the MSM weights.

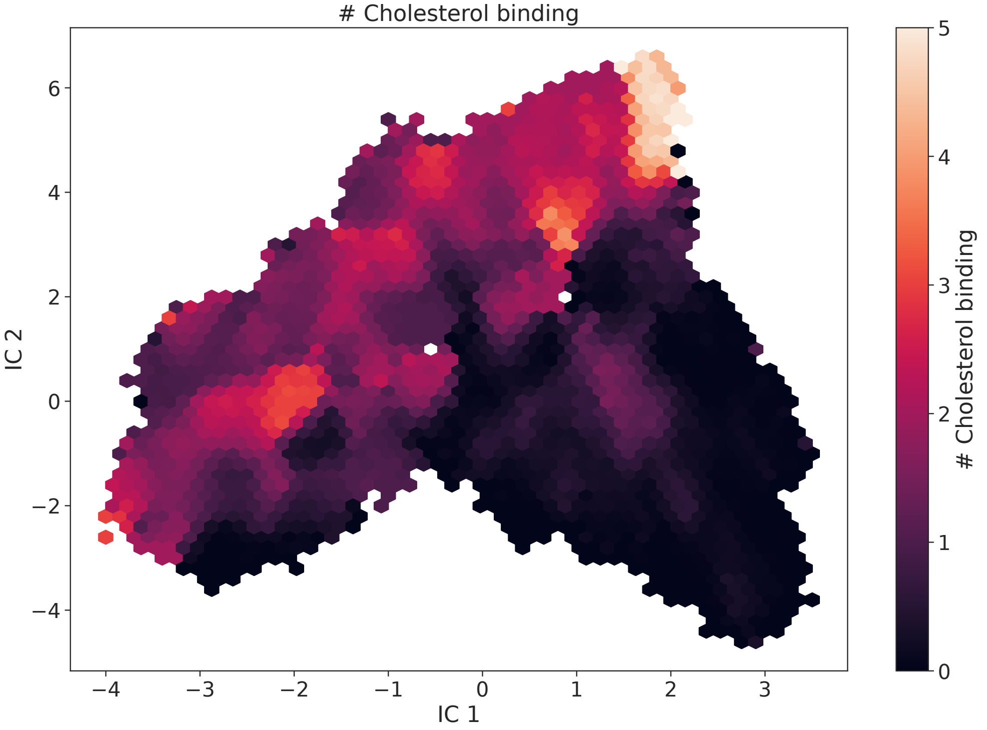


**Figure S7. Number of cholesterol molecules bound to the intersubunit binding site mapped onto IC1-IC2 coordinates.**

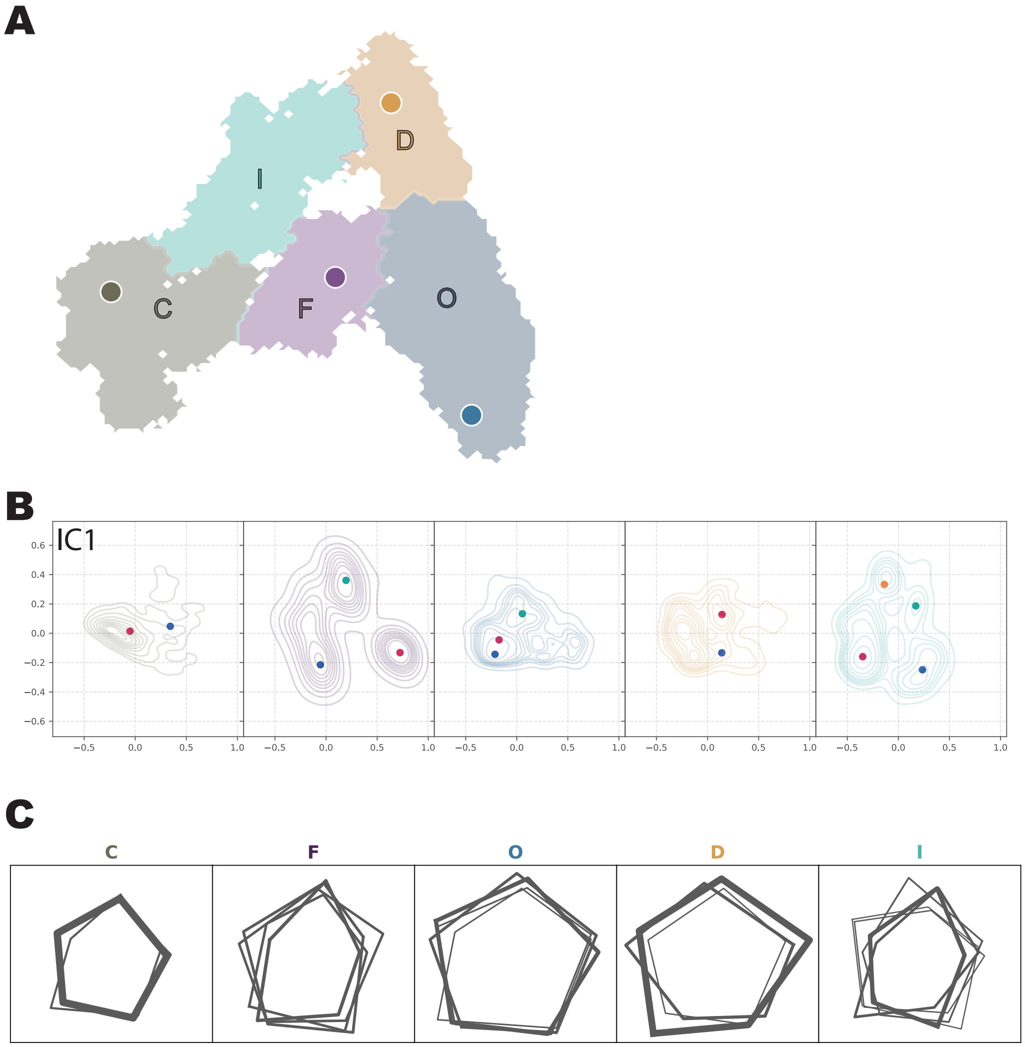


**Figure S8. MSM mapping and asymmetric analysis.**(**A**) Mapping of a coarse-grained MSM with five macrostates onto IC1-IC2 coordinates in the CHOL system. Four structural models are projected onto the plot. (**B**) Symmetry-aware multidimensional scaling (MDS) projected sub Independent Components (IC1) onto the MDS1-MDS2 space. The zero point represents perfect symmetry with mean value, while a shifted distribution suggests sampling of asymmetric conformations. Representative snapshots in the free-energy basins are shown as dots (**C**) Representative snapshots of the 9′ $C_\alpha$ pentagon captured in each macrostate at each local free energy minimum. The thickness of each pentagon indicates the relative population of that conformation within the corresponding macrostate.

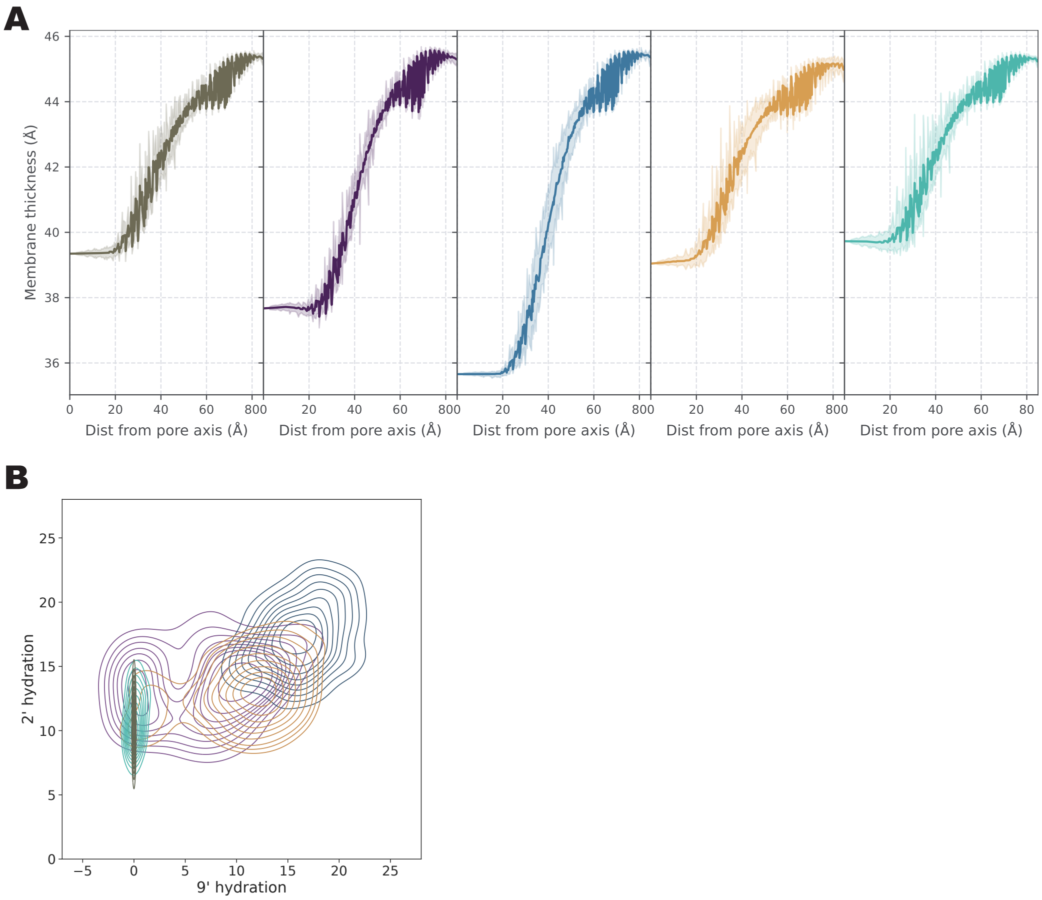


**Figure S9. Membrane compression and pore hydration profile in the CHOL system.** (**A**) Membrane thickness profiles in each macrostate, obtained by radially averaging the membrane thickness centered at the pore axis. (**B**) Histogram of the number of water molecules around the 9′ residue versus the 2′ residue. This analysis was performed within a cylindrical region centered at each residue, extending ±2 Å along the pore axis.

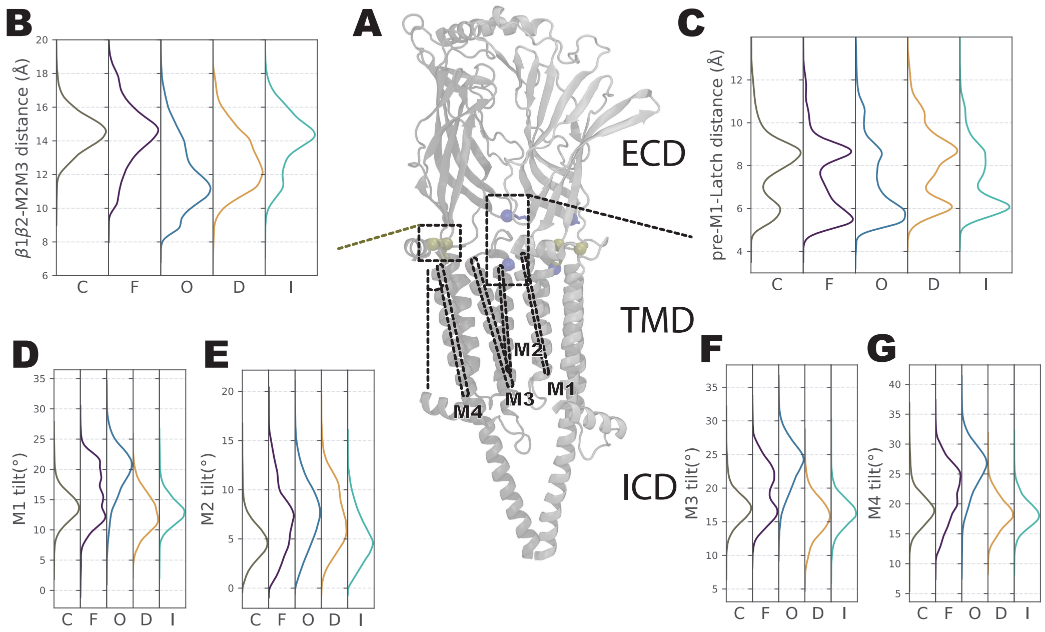


**Figure S10. Sequential conformational changes of coupling region and transmembrane domain in the CHOL system.** (**A**) Visualization of two consecutive subunits, highlighting the contact between the pre-M1 loop and latch helix (shown in tan) and the contact between the β1-β2 loop and M2-M3 loop (shown in blue). The M1, M2, M3, and M4 helices are labeled with dashed lines. (**B**) Histogram showing the distribution of contacts between the pre-M1 loop and latch helix for each macrostate. (**C**) Histogram showing the distribution of contacts between the β1-β2 loop and M2-M3 loop for each macrostate. (**D, E, F, G**) Histograms showing the distribution of tilt angles for the M1, M2, M3, and M4 helices, respectively, in each macrostate. All distributions are weighted by the MSM weights.

### Appendix 1

#### Supplementary Note: Symmetric Toy model

Synthetic timeseries data generated from a given markovian transition matrix between global states. Eight for a dimer with three sub-states (***Figure 1***); another eight for a trimer with two sub-states (***Figure 2***); 16 for a tetramer with two sub-states (***Figure 3***); In total six features comprised three observables in each subsystem. Conventional time-lagged independent component analysis (TICA) resolves all timescales while SymTICA (correctly) only resolves the degenerate model for the dynamics.

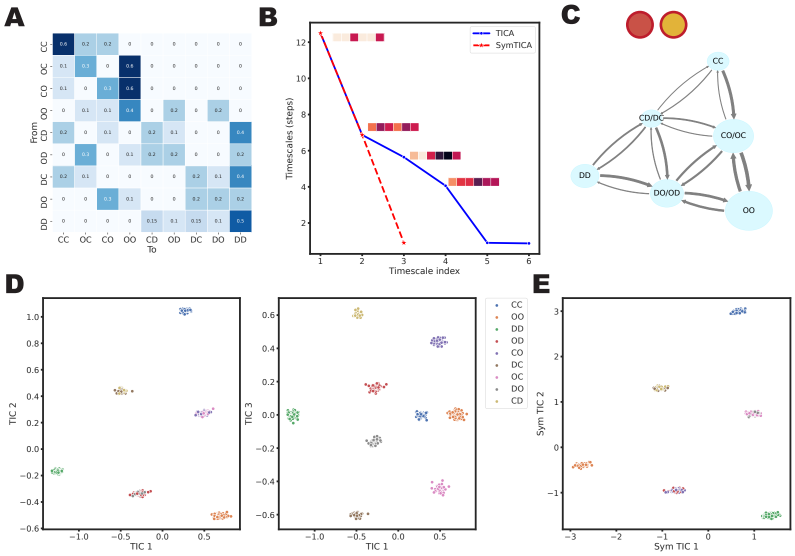


**Appendix 1—figure 1. Three-substate dimer toy model.** (**A**) Transition probability matrix between all global states that comprise ordered subsystem states. (**B**) Resolved timescales by TICA and SymTICA. The schematic representation of projection vectors for each dimension is shown. SymTICA only resolves projections that are identical for each subsystem. (**C**) The degenerate Markov state diagram in which intermediate states are lumped together. (**D**) Data projected onto TIC 1-2-3 space by TICA, colored by their true global state. (**E**) Data projected onto Sym TIC 1-2 space by SymTICA.

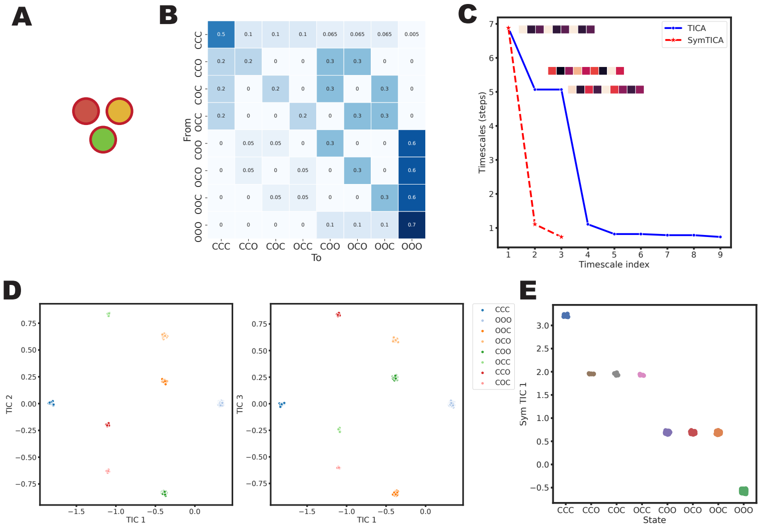

**Appendix 1—figure 2. Two-substate trimer toy model.** (**A**) Schematic image of a trimer toy model. (**B**) Transition probability matrix between all global states that comprise ordered subsystem states. (**C**) Timescales resolved by TICA and SymTICA. The schematic representation of projection vectors for each dimension is shown. (**D**) Data projected onto TIC 1-2-3 space by TICA, colored by their true global state. (**E**) Data projected onto Sym TIC 1 space by SymTICA.

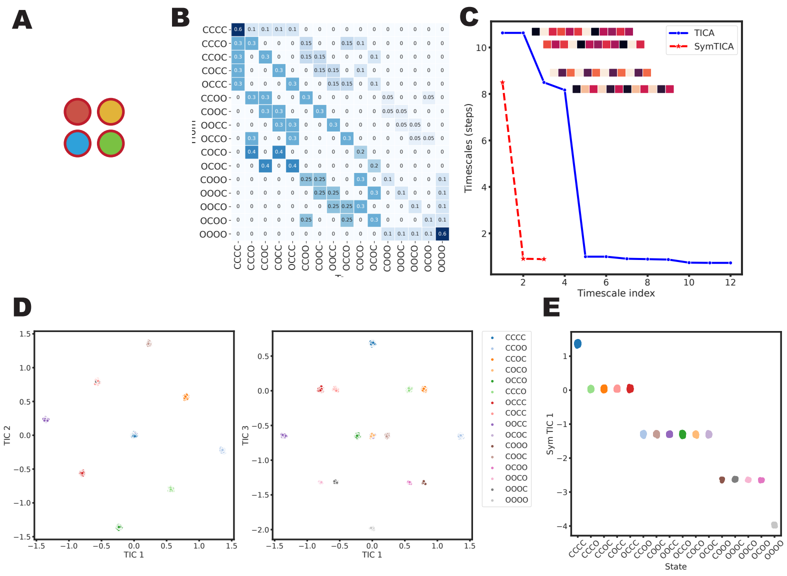


**Appendix 1—figure 3. Two-substate tetramer toy model** (**A**) Schematic image of a tetramer toy model. (**B**) Transition probability matrix between all global states that comprise ordered subsystem states. (**C**) Timescales resolved by TICA and SymTICA. The schematic representation of projection vectors for each dimension is shown. (**D**) Data projected onto TIC 1-2-3 space by TICA, colored by their true global state. (**E**) Data projected onto Sym TIC 1 space by SymTICA.
